# ClassificaIO: machine learning for classification graphical user interface

**DOI:** 10.1101/240184

**Authors:** Raeuf Roushangar, George I. Mias

## Abstract

Machine learning methods are being used routinely by scientists in many research areas, typically requiring significant statistical and programing knowledge. Here we present ClassificaIO, an open-source Python graphical user interface for machine learning classification for the scikit-learn Python library. ClassificaIO provides an interactive way to train, validate, and test data on a range of classification algorithms. The software enables fast comparisons within and across classifiers, and facilitates uploading and exporting of trained models, and both validation and testing data results. ClassificaIO aims to provide not only a research utility, but also an educational tool that can enable biomedical and other researchers with minimal machine learning background to apply machine learning algorithms to their research in an interactive point-and-click way. The ClassificaIO package is available for download and installation through the Python Package Index (PyPI) (http://pypi.python.org/pypi/ClassificaIO) and it can be deployed using the “import” function in Python once the package is installed. The application is distributed under an MIT license and the source code is publicly available for download (for Mac OS X, Linux and Microsoft Windows) through PyPI and GitHub (http://github.com/gmiaslab/ClassificaIO, and https://doi.org/10.5281/zenodo.1320465).

## Introduction

Recent advances in high-throughput technologies, especially in genomics, have led to an explosion of large-scale structured data (e.g. RNA-sequencing and microarray data) (1). Machine learning methods (classification, regression, clustering, etc.) are routinely used in mining such big data to extract biological insights in a range of areas within genetics and genomics (2). For example, using unsupervised machine learning classification methods to predict the sex of gene expression microarrays donor samples (3), using genome sequencing data to train machine learning models to identify transcription start sites (4), splice sites (5), transcriptional promoters and enhancers regions (6). Recent examples of using machine learning classification methods include their use to detect neurofibromin 1 tumor suppressor gene inactivation in glioblastoma (7), and to identify reliable gene markers for drug sensitivity in acute myeloid leukemia (8). Many advanced machine learning algorithms have been developed in the recent years. Scikit-learn (9) is one of the most popular machine learning libraries in Python with a plethora of thoroughly tested and well-maintained machine learning algorithms. However, these algorithms are primarily aimed at users with computational and statistical backgrounds, which may discourage many biologists, biomedical scientists or beginning students (who may have minimal machine learning background but still want to explore its application in their research) from using machine learning. Ching et al (2018) recently highlighted the role of deep learning (a class of machine learning algorithms) currently plays in biology, and how such algorithms present new opportunities and obstacles for a data-rich field such as biology (10).

Several open source machine learning applications, such as KNIME (11) and Weka (12) written in Java and Orange (13) written in Python, have been developed with graphical user interfaces. The dataflow process for most of these applications is generally graphically constructed by the user, in the form of placing and connecting widgets by drag-and-drop. Such graphical workflow and representation of data input, processing and output is visually appealing, but can be computationally demanding (memory, storage, processing, etc.) and limiting in algorithm comparison, since each machine learning algorithm can have many different parameters. These tools are very mature with numerous algorithms, and well documented. However, they can be intimidating for machine learning beginners and students that want to preform simple tasks such as data classification. Also, scikit-learn has comprehensive documentation (14), and many online resources, including though Kaggle (15) and Stack Overflow (16), and a large online user base, which make scikit-learn a very popular package for machine learning beginners learning using Python.

Here, we present ClassificaIO, an open-source Python graphical user interface (GUI) for supervised machine learning classification for the scikit-learn library. To the best of our knowledge, no standalone GUI exists for the scikit-learn library. ClassificaIO’s core aim is to provide a, machine learning research, teaching and educational tool that is visually minimalistic and computationally light interactive interface, that can give access to a range of state-of-the-art classification algorithms to machine learning beginners with some basic knowledge of Python and using a terminal, and with broad background in machine learning, allowing them to use machine learning and apply it to their research. What distinguishes ClassificaIO from other similar applications is:

- Cross-platform implementation for Mac OS X, Linux, and Windows operating systems
- Interactive point-and-click GUI to 25 supervised classification algorithms in scikitlearn
- Accessible clickable links, to scikit-learn’s well-written online documentation for each implemented classification algorithms
- Simple upload of all data files with dedicated buttons; with robust CSV reader, and a displayed history-log to track uploaded files, files names and directories
- Fast comparisons within and across classifiers to train, validate, and test data
- Upload and export of ClassificaIO trained models (for future of a trained model without the need to retrain), and export of both validated, and tested data results
- Small application footprint in terms of disk space usage (<2 MB)

## Materials and methods

### ClassificaIO implementation

ClassificaIO has been developed using the standard Python interface Tkinter module to the Tk GUI toolkit (17), for Mac OS X (using High Sierra ≥ 10.13), Linux (using Ubuntu 18.04-64 bit), and Windows (using Windows 10 64-bit), Table 1. It uses external packages including: Tkinter, Pillow, Pandas (18), NumPy (19), scikit-learn and SciPy (20). To avoid any system errors, crashes, and crude fonts, we recommend not to install ClassificaIO using integrated environment package installers – instead, native installation of ClassificaIO and dependencies (using pip for Mac and Windows, and pip3 and apt-get for Linux) is encouraged. Once installed, ClassificaIO can be deployed using the ‘import’ function. A ClassificaIO user manual uploaded as supplementary file S1_Manual (hereafter referred to as S1) is also distributed with ClassificaIO GUI and can be accessed directly through the ‘HELP’ button at the upper left of the GUI, that points the user’s default browser to ClassificaIO’s online user manual on GitHub. Some basic knowledge of Python and accessing it through a terminal are required for installation and running the software. Additional ClassificaIO software information is provided in Table 1.

**Table 1.**
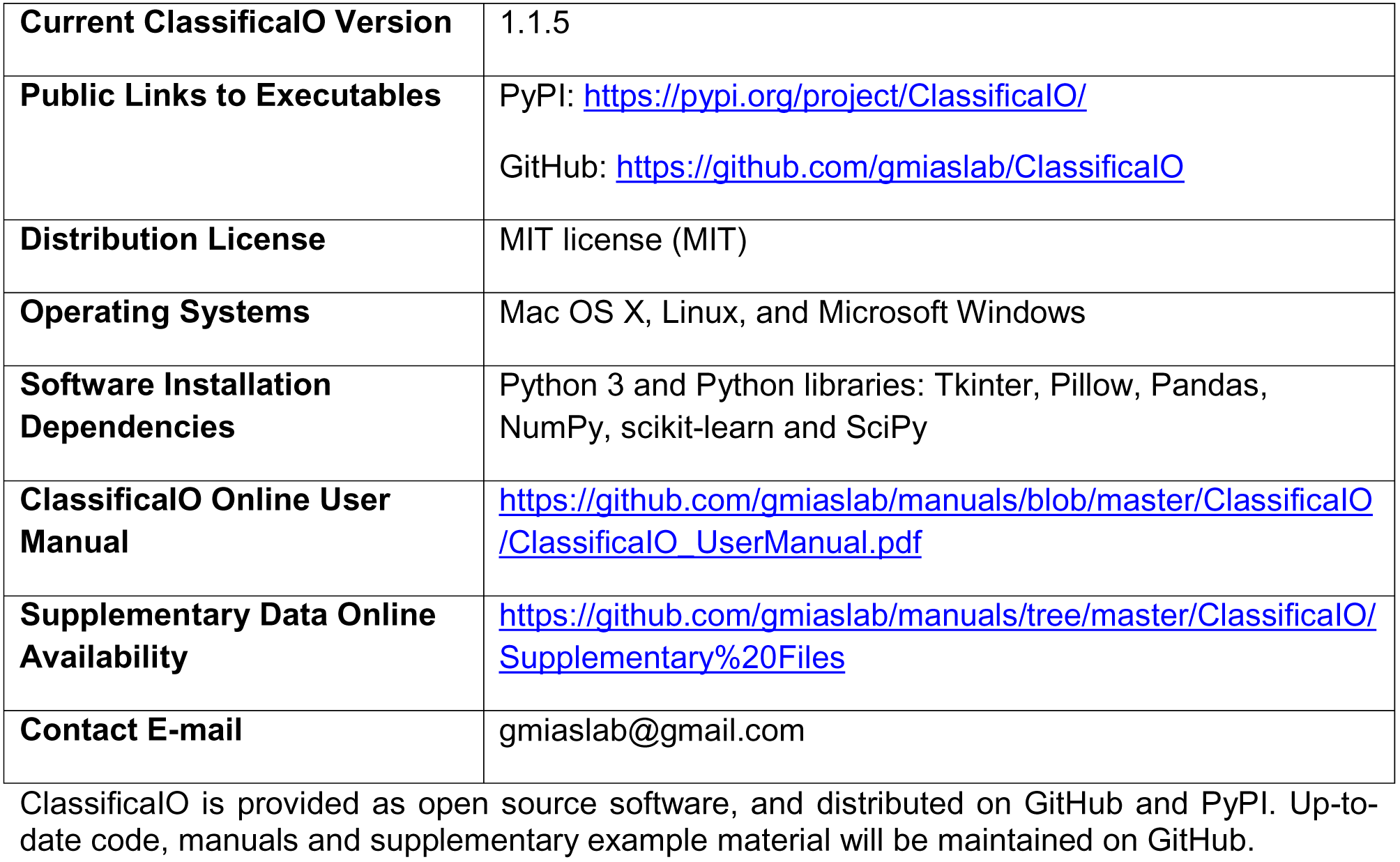
**ClassificaIO software information**

### ClassificaIO backend

ClassificaIO implements 25 scikit-learn classification algorithms for supervised machine learning. A list of all these algorithms, their corresponding scikit-learn functions, and immutable (unchangeable) parameters with their default values are presented in S1 Table 1, and ClassificaIO’s workflow is outlined in Fig 1. Once training and testing data are uploaded to the front-end as described below, a classifier selection is made and submitted, ClassificaIO’s backend calls the scikit-learn selected classifier, including any values from manually set parameters to create the model. Otherwise, the default parameters values are used instead. For example, for “LogisticRegression”, the model is defined in the scikit-learn library as a class, in terms of Python code used in the backend, the details are outlined in the scikit-learn documentation:

*sklearn.linear_model.LogisticRegression(penalty=‘l2’, dual=False, tol=0.0001, C=1.0, fit _intercept=True, intercept_scaling=1, class_weight=None, random_state=None, solver= ‘liblinear’, max_iter=100, multi_class=‘ovr’, verbose=0, warm_start=False, n_jobs=1)*

**Fig 1.**
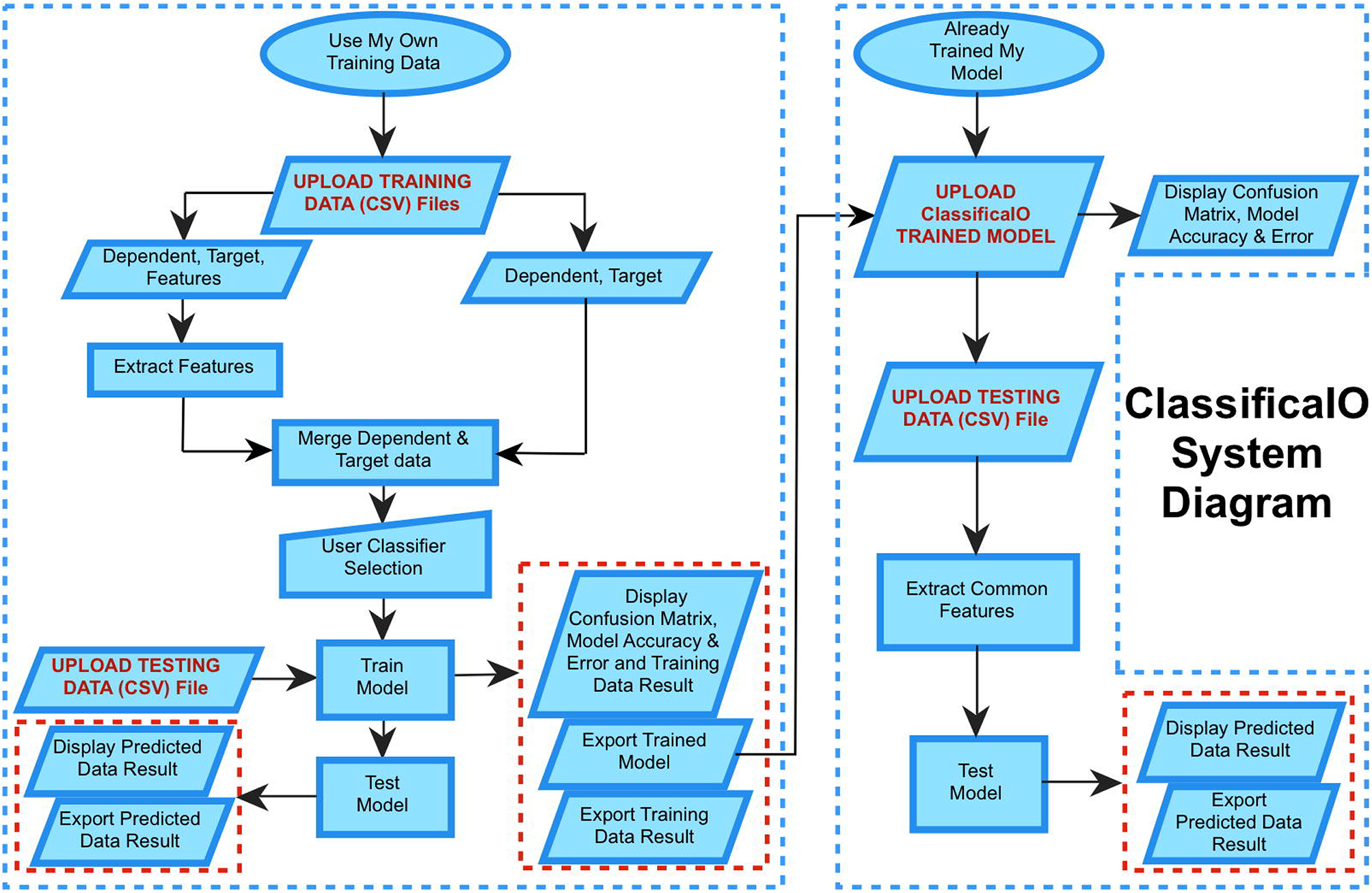
**ClassificaIO workflow.** The diagram summarizes the graphical user interface and backend functionality/workflow for ClassificaIO Use My Own Training Data and Already Trained My Model windows.

The inputs to the class, within the parentheses, such as “penalty”, “dual”, “tol”, etc., correspond to the model parameters, followed by an equal sign assigning the default values for these parameters. Rather than typing the values, the ClassificaIO GUI displays these parameters with input fields and radio buttons, for each classifier, initially populated by the default values. More information is available for all the parameters in the GUI, through a link for each classifier in the interface named “Learn More”. The link directs the default browser to the scikit-learn online documentation of the selected classifier, and connects to the underlying backend documentation, and online parameter descriptions. The details and code complexity of the backend implementation are effectively hidden from the user, who can interact with the ClassificaIO GUI to set the relevant parameters, or leave them unchanged as default values. On the training data, ClassificaIO fits the estimator for classification using the scikit-learn ‘fit’ method, e.g. fit(x_train, y_train), to train (learn from the model), and uses the scikit-learn ‘predict’ method, e.g. predict(x_validation), to validate the model. Finally, ClassificaIO predicts new values using the scikit-learn ‘predict’ method again but on the testing data, e.g. predict(testing_X), for implementing the model on new data that have not been used in model training.

### ClassificaIO functionalities

ClassificaIO’s GUI consists of three windows: ‘Main’, ‘Use My Own Training Data’, and ‘Already Trained My Model’. Each window is actually implemented within the code as a class with several functions/methods that are dynamically connected to provide the GUI. ClassificaIO’s Main window, Fig 2, has two buttons: (i) the ‘Use My Own Training Data’ button, which when clicked allows the user to train and test classifiers using their own training and testing data, (ii) the ‘Already Trained My Model’ button, which when clicked allows the user to use their own already ClassificaIO trained model and testing data.

**Fig 1.**
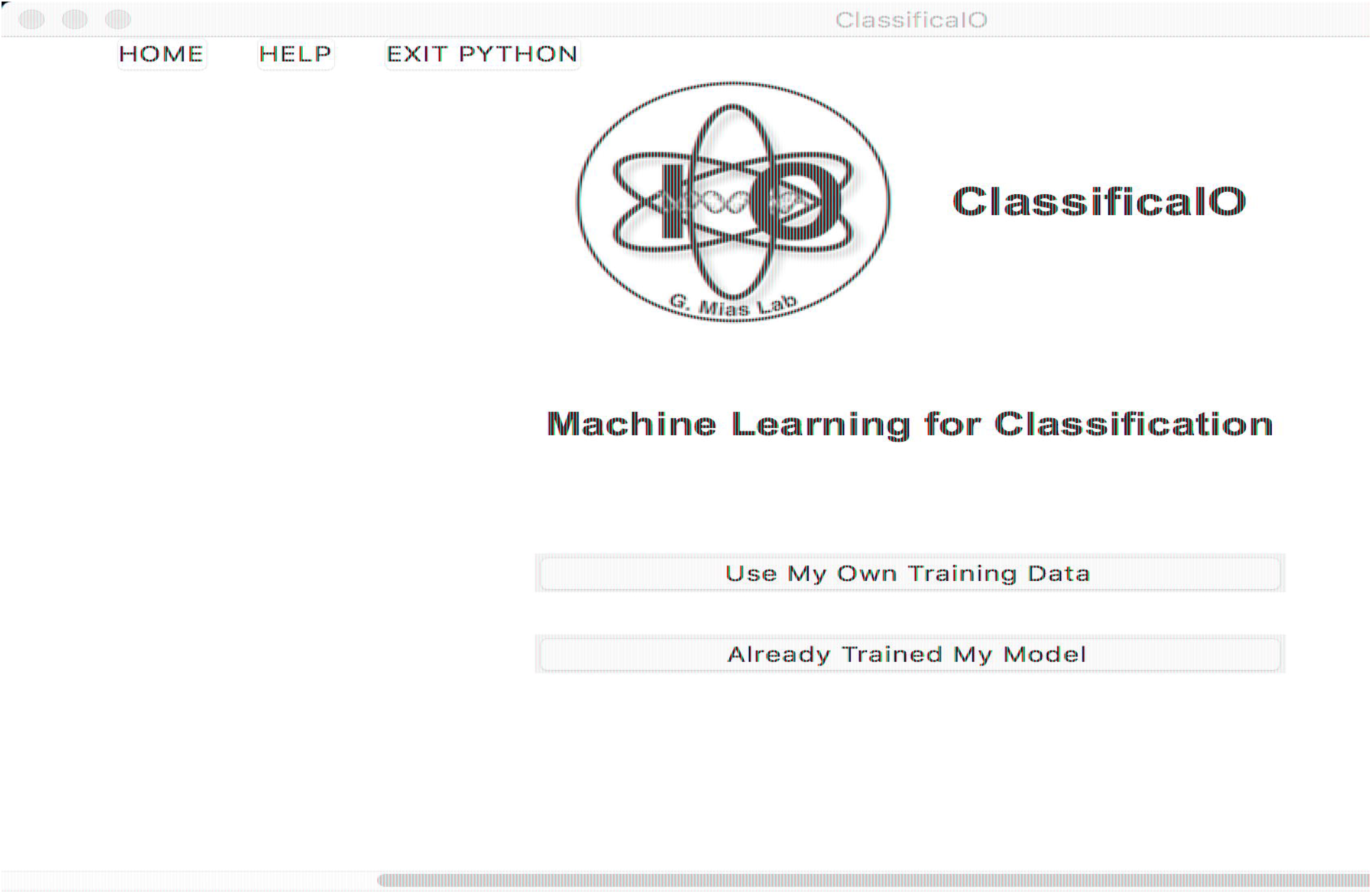
**ClassificaIO main window.** The main window appears on the screen after typing ‘ClassificaIO.gui()’ in a terminal or a Python interpreter.

### Data input

For the ‘Use My Own Training Data’ window, Fig 3a, by clicking the corresponding buttons in the ‘UPLOAD TRAINING DATA FILES’ and ‘UPLOAD TESTING DATA FILE’ panels, a file selector directs the user to upload all required comma-separated values (CSV) data files (‘Dependent and Target’ or ‘Dependent, Target and Features’ and ‘Testing Data’) (S1 Fig 1). Briefly, the dependent data represent the data on which the model will depend on for learning, and the target data is the annotation, i.e. what is going to be predicted. The dependent data have attributes (also known as features) that take values (measurements/results) for each contained object (i.e. each sample). Further details on data files formats and examples are provided in the S1 Figs 3(a, b) and S2-S5. A history of all uploaded data files (file name and directory) is automatically saved in the ‘CURRENT DATA UPLOAD’ panel (S1 Figs 2 and 7(c)).

**Fig 3.**
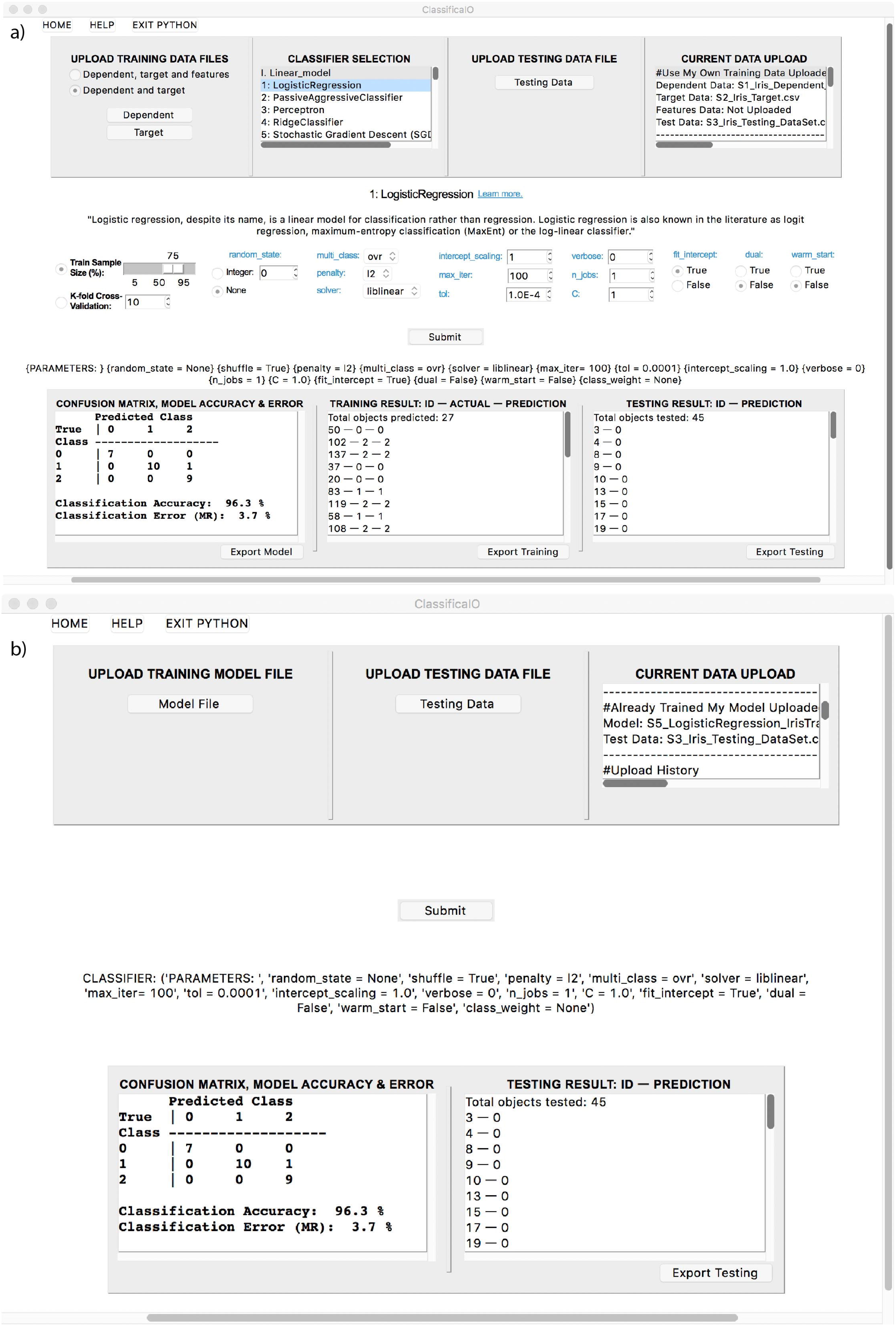
**ClassificaIO user interface (Mac OS shown).** As described in ClassificaIO implementation section, (a) an example ‘Use My Own Training Data’ window with uploaded training and testing data files, selected logistic regression classifier, populated classifier parameters, and output classification results. (b) A corresponding ‘Already Trained My Model’ window with uploaded logistic regression ClassificaIO trained model and testing data file, and output classification result.

### Classifier selection

After the data is uploaded, the user can select between all 25 different widely used classification algorithms (including logistic regression, perceptron, support vector machines, k-nearest neighbors, decision tree, random forest, neural network multi-layer perceptron, and more). The algorithms are integrated from the scikit-learn library, and allow the user to train and test models using their own uploaded data. Each classifier can be easily selected by clicking the corresponding classifier name in the ‘CLASSIFER SELECTION’ panel (see S1 ‘Classifier selection’ section). Once classifier selection is completed, a brief description for the classifier with an underlined clickable link that reads “Learn more” right next to the classifier name (S1 Fig 4(a)) and the classifier parameters will populate (S1 Fig 4(c)). If “Learn more” is clicked, the link directs the default web browser to open scikit-learn’s online well-written documentation that explains the specific classifier parameters, with explanation for each parameter and its use, and how to tune/optimize each parameter to get the best performance. ClassificaIO provides the user with an interactive point-and-click interface to set, modify, and test the influence of each parameter on their data. The user can switch between classifiers and parameters through point-and-click, which enables fast comparisons within and across classifier models.

### Model training

Both train-validate split and cross-validation methods (which are necessary to prevent/minimize overfitting) will populate with each classifier that can be used for data training (S1 Fig 4(b)). Training, validating and testing are all performed after pressing the submit button (middle of Fig. 3(a)).

### Results output

After model training and testing is completed, the confusion matrix, classifier accuracy and error are displayed in the ‘CONFUSION MATRIX, MODEL ACCURACY & ERROR’ panel, bottom of Fig 3(a) and S1 Fig 5(a). Model validation data results are displayed in the ‘TRAINING RESULT: ID – ACTUAL – PREDICTION’ panel, S1 Fig 5(b), and testing data results are displayed in the ‘TESTING RESULT: ID – PREDICTION’ panel, S1 Fig 6(b).

### Model export

By clicking on the ‘Export Model’ button (see Fig 3(a) bottom left and S1 Fig 5(a)), the user can export trained models to save for future use without having to retrain. A previously exported ClassificaIO model can then be used for testing of new data in the ‘Already Trained My Model’ window, Fig 3(b), by clicking the ‘Model file’ button in the ‘UPLOAD TRAINING MODEL FILE’ panel (S1 Fig 7(a)).

### Results export

Full results (trained models, both validated and tested data, and uploaded files names and directories) for both windows, Fig 3(a and b), can be exported as CSV files for further analysis for publication, sharing, or later use (for more details on the exported trained model and data file formats, see S7 and S8).

### Results: Illustrative examples and data used

To illustrate the use of the interface and classification, we have used in this manuscript the following two examples (also see the S1 for more details).

i. **Iris prediction using Iris dataset.** To demonstrate the interface and classification, we used the so-called Fisher/Anderson iris dataset (21, 22). This dataset is used widely as a prototype to illustrate classification algorithms, not only of biological data but in general machine learning implementations. The dataset consists of fifty samples each for three different species of iris flowers (Setosa, Versicolor and Viginica), with sepal length and width, and petal length and width provided as measurements. For more details on the iris data files format, see S1 Fig 3(a,b) and S2-S5.
ii. **Sex prediction using microarray gene expression data.** In this example, provided in ClassificaIO user manual (S1), we used raw microarray gene expression data, from Gene Expression Omnibus (GEO) (23) to predict each sample donor’s sex. This is often necessary in metadata analyses, using publicly available gene expression datasets for reanalysis, as samples annotations on GEO may be missing information, including sample donor’s sex. To illustrate the classification/sex prediction we used two datasets, GSE99039 (24) (training data) and GSE18781 (25) (testing data). In both GSE99039 and GSE18781 datasets, we used 121 and 25 samples respectively, for which RNA from peripheral blood mononuclear cells was assayed using Affymetrix Human Genome U133 Plus 2.0 Array (accession GPL570). The Y chromosome gene expression values were used in ClassificaIO as training and testing data to predict samples donor’s sex. Using the ‘Linear SVC’ model with “k-fold cross validation” (10-fold), resulted into a model with 99% accuracy for sample donor’s sex prediction (in the displayed example). For more details on the pre-processing of the raw gene expression data, files format, and Y chromosome probes ids, and final result, see S1 ‘Additional Examples: Ex2’ section and Figs 9 and 10.

## Discussion

We have presented ClassificaIO, a GUI that implements the scikit-learn supervised machine learning classification algorithms. The scikit-learn package is one of the most popular in Python with well-written documentation, and many of its machine leaning algorithms are currently used for analyzing large and complex data sets in genomics. Our interface aims to provide an interactive machine learning research, teaching and educational tool to do machine learning analysis without the requirement of advanced computational and machine learning knowledge using scikit-learn. ClassificaIO is provided as an open source software, and its back-end classes and functions allow for rapid development. We anticipate further development, aided by the scikit-learn library developer community to integrate additional classification algorithms, and extend ClassificaIO to include other machine leaning methods such as regression, clustering, and anomaly detection, to name but a few.

## Acknowledgements

None

## Supporting information

**S1 Fig 1. Graphical control element dialog box.** (a) Dependent data file selected for upload. (b) Selected target data file to upload. N.B. each file selection has to be done one at a time.

**S1 Fig 2. Current data upload panel.** Both dependent and target data file names shown (red boxes). Scroll down for uploaded data files directories.

**S1 Fig 3(a) Dependent data.** Example of partial dependent data file format. Testing data (not shown) uses the same format.

**S1 Fig 3(b) Target data.** Example of partial target data file format where the targets correspond to setosa = 0, versicolor = 1, and virginica = 2. Versicolor and virginica are not visible in this screenshot.

**S1 Fig 4. Selected logistic regression classifier.** The interface for each selected classifier, has uniform features. (a) Classifier definition is displayed, together with an underlined clickable link that reads “Learn more” next to the classifier name. (b) Training methods with ‘Train Sample Size (%)’ method selected. (c) The classifier parameters set to their default values.

**S1 Fig 5. Trained logistic regression classifier. (a)** Trained model using 78 data points (75% of 105 data points), classifier evaluation (confusion matrix, model accuracy and error). (b) Model validated using 27 data points (25% of 105 data points).

**S1 Fig 6. Tested logistic regression classifier.** (a) Upload testing data panel. (b) Model tested using 45 data points.

**S1 Fig 7. ‘Already Trained My Model’ window.** (a) Upload ClassificaIO trained model panel. (b) Upload testing data panel. (c) Current data upload panel with both model and testing data files names shown (red boxes). (d) Model preset parameters. (e) Trained model result and model evaluation (confusion matrix, model accuracy and error). (f) Model testing result.

**S1 Fig 8. Training and testing using gene expression data.** (a) selected k-nearest neighbors classifier with trained and tested the data using the default parameters values, (b) Same classifier selected with trained and tested data but using different parameters values.

**S1 Fig 9. Trained linear support vector machine classifier.** Trained model using GSE99039 121 data points and k-fold cross validation, classifier evaluation (confusion matrix, model accuracy and error). Model validated and tested model using GSE18781 25 data points.

**S1 Fig 10. Features data.** Example of partial features data file format where each Affymetrix probe id correspond to a Y chromosome gene.

**S1_Manual. ClassificaIO user manual.** User manual for ClassificaIO, including all supplementary Figs 1-10 as well as working examples implementing using supplementary files S2-S8.

**S2_Iris_Dependent_DataSet.csv. Iris dependent data set (105 data points).**

**S3_Iris_Target.csv. Iris Target data set (105 labels).**

**S4_Iris_Testing_DataSet.csv. Iris testing data set (45 data points).**

**S5_Iris_FeatureNames.csv. Example Iris features (2 features: sepal length and petal width).**

**S6_LogisticRegression_IrisTrainedModel.pkl. Example ClassificaIO trained model using logistic regression.**

**S7_IrisTrainValidationResult.csv. Example ClassificaIO testing result using logistic regression.**

**S8_IrisTestingResult.csv. Example ClassificaIO validation result using logistic regression.**

